# A retrospective analysis of gender parity in scientific authorship in a biomedical research centre

**DOI:** 10.1101/2020.02.24.962175

**Authors:** Rinita Dam, Syed Ghulam Sarwar Shah, Maria Julia Milano, Laurel D Edmunds, Lorna R Henderson, Catherine R Hartley, Owen Coxall, Pavel V Ovseiko, Alastair M Buchan, Vasiliki Kiparoglou

## Abstract

**Objective:** Scientific authorship is a vital marker of success in academic careers and gender equity is a key performance metric in research. However, there is little understanding of gender equity in publications in biomedical research centres funded by the National Institute for Health Research (NIHR). This study assesses the gender parity in scientific authorship of biomedical research.

**Design:** A retrospective descriptive study.

**Setting:** NIHR Oxford Biomedical Research Centre.

**Data:** 2409 publications accepted or published from 1 April 2012 to 31 March 2017.

**Main outcome measures:** Gender of authors, defined as a binary variable comprising either male or female categories, in six authorship categories: first author, joint first authors, first corresponding author, joint corresponding authors, last author and joint last authors.

**Results:** Publications comprised clinical research (39%, n=939), basic research (27%, n=643), and other types of research (34%, n=827). The proportion of female authors as first author (41%), first corresponding authors (34%) and last author (23%) was statistically significantly lower than male authors in these authorship categories. Of total joint first authors (n=458), joint corresponding authors (n=169), and joint last authors (n=229), female only authors comprised statistically significant smaller proportions i.e. 15% (n=69), 29% (n=49) and 10% (n=23) respectively, compared to male only authors in these joint authorship categories. There was a statistically significant association between gender of the last author(s) with gender of the first author(s) (χ ^2^ 33.742, P < 0.001), corresponding author(s) (χ^2^ 540.774, P < 0.001) and joint last author(s) (χ ^2^ 91.291, P < 0.001).

**Conclusions:** Although there are increasing trends of female authors as first authors (41%) and last authors (23%), female authors are underrepresented compared to male authors in all six categories of scientific authorship in biomedical research. Further research is needed to encourage gender parity in different categories of scientific authorship.

**Strengths and limitations of this study:**

- This is the first study to investigate gender parity in six categories of scientific authorship: first authors, first corresponding authors, last authors and three joint authorship categories i.e. joint first authors, joint corresponding authors and joint last authors in biomedical research.
- This study provides an important benchmark on gender equity in scientific authorship for other NIHR funded centres and organisations in England.
- The generalisability of the findings of this study may be limited due to differences in medical specialities, research areas, institutional cultures, and levels of support to individual researchers.
- Using secondary sources for determining the gender of authors may have limitations, which could be avoided by seeking relevant information from original authors and institution affiliation at the time of submission.

## INTRODUCTION

Promoting Responsible Research and Innovation (RRI) is a major strategy of the “Science with and for Society” work programme of the European Union’s (EU) Horizon 2020 Framework Programme for Research and Innovation[1]. RRI aims to build capacity and develop innovative ways to connect science to society[2]. The RRI approach enables all societal members (such as researchers, citizens, policymakers, businesses and third sector organisations) to work together during the research and innovation process in order to better align research and innovation with the values, needs and expectations of the society[1,2]. The RRI strategy includes the “keys” of public engagement, open access, gender, ethics and science education, and two further “keys”: sustainability and social justice, which have been added recently [3]. The idea is that by prioritising these key components of RRI, it would help make science more attractive to young people and society, and raise awareness of the meaning of responsible science[2].

We have focussed on the ‘gender’ element of the RRI because it is imperative to advance gender equality within research institutions, as well as within the design and content of research and innovation[1]. The issue of enhancing female participation in economic decision-making has become prominent in the national, European and international spheres, with a particular focus on the economic dimension of gender diversity[4]. In order to achieve a fair female participation within positions of power, it is recommended that women should hold half of the total seats in board rooms[5], however, a ratio between 40% and 60%, also known as a “gender balance zone”[6], is considered acceptable – a threshold that is set by the European Commission[4].

From the perspective of gender equality in academia and scientific research, gender parity in scientific authorship is an important measure of achievement. The term gender parity refers to “the equal contribution of women and men to different dimensions of life” and it is operationalised as a “relative equality in terms of numbers and proportions of women and men” for a particular indicator”[7]. Gender (dis)parity in scientific authorship has important implications for gender equity in academic advancement[8] because scientific authorship is commonly used as a measure of academic productivity that is used for performance management, reward, and recognition[9,10]. The acceleration of women’s advancement and leadership in research is one of the stated objectives of the National Institute of Health Research (NIHR) in the United Kingdom (UK) and it is imperative for the RRI in the wider European research area. Yet, there is limited research concerning gender equity in scientific authorship of translational research funded through NIHR biomedical research centres (BRCs).

In the UK, women currently outnumber men in medical schools[11], however, a persistent gender disparity in scientific publications remains[10,12–23]. While the proportion of women as first and senior authors of original medical research has increased over the past few decades[24], women are still significantly underrepresented as authors of research articles in medical journals, especially as first and senior authors[14,22,23,25]. For example, in radiology the proportion of women as first author increased from 8% in 1978 to 32% in 2013 and senior author increased from 7% in 1978 to 22% in 2013[23]. Similarly, in gastroenterology the proportion of women as first author increased from 9% in 1992 to 29% in 2012, and senior author increased from 5% in 1992 to 15% in 2012[14].

The profile of gender equity in higher education and research has been raised by the introduction of Athena SWAN-linked funding incentives by the NIHR[26–28]. While Athena SWAN awards are useful markers of achievement for higher education institutions and research institutes, they alone are insufficient to assess and monitor the progress of NIHR BRCs towards gender equity[29]. Currently, the proportion of women and the rate of their achievements are not tracked routinely by the NIHR BRCs and little is known about how much women contribute to scientific research and innovation in the BRCs. It is important to inform the acceleration of women’s advancement and leadership in translational research in line with the stated objectives of the NIHR within the UK and RRI within the wider European research area through the collection of gender-disaggregated bibliometric data and analysis of scientific authorship by gender.

For addressing the paucity of empirical research on women’s advancement and leadership in translational research in the UK and Europe, a recent study on gender equity in Neurology suggests the need for institutions to take a systematic approach to addressing gender disparities that involve customised, defined metrics and transparent reporting to stakeholders[30].

The aim of this study was to assess the gender parity in six types of scientific authorship in biomedical research.

## METHODS

### Study design

A retrospective descriptive study.

### Setting

This study was conducted at the NIHR Oxford BRC, which is research collaboration between the Oxford University Hospitals NHS Foundation Trust and the University of Oxford[31]. The aim of NIHR BRCs is to support translational research and innovation to improve healthcare for patients[32]. During the study period (April 2012-March 2017), the NIHR Oxford BRC was awarded £96m to support research across nine research themes, five cross-cutting themes, and a range of underpinning platforms. The research themes included Blood, Cancer, Cardiovascular, Dementia and Cerebrovascular Disease, Diabetes, Functional Neuroscience and Imaging, Infection, Translational Physiology, and Vaccines. The crosscutting themes included Genomic Medicine, Immunity and Inflammation, Surgical Innovation and Evaluation, Biomedical Informatics and Technology, and Prevention and Population Care. The major underpinning platforms included a Biorepository, Education and Training, Public Engagement, and Research Governance. It is a contractual requirement to report the number of BRC supported publications published by researchers funded or supported by the NIHR research funds on an annual basis. Additionally, the NIHR uses bibliometric analyses to inform eligibility for NIHR funding[33,34]. This study was carried out as part of a wider programme of research on the markers of achievement for assessing and monitoring gender equity in translational research organisations[29].

### Data

Data comprised translational research publications published by researchers funded or supported by the NIHR Oxford BRC. The eligibility criteria for inclusion of a publication were funding or support by the NIHR Oxford BRC and publication between April 2012 and March 2017. Based on these criteria, 2409 publications were identified. These publications were classified as: basic science studies, clinical studies (both trial and non-trial studies), and other studies (comments, editorials, systematic reviews, reviews, opinions, meeting reports, guidelines and protocols).

### Main outcome measures

The main outcome measures were: (1) Gender of authors, defined as a binary variable comprising, either male or female categories, (2) Six categories of scientific authorship: first author, joint first authors, first corresponding author, joint corresponding authors, last author and joint last authors (Figure 1). These categories are conventionally associated with the highest amount of contribution, credit and prestige[10,17].

**Figure 1.**
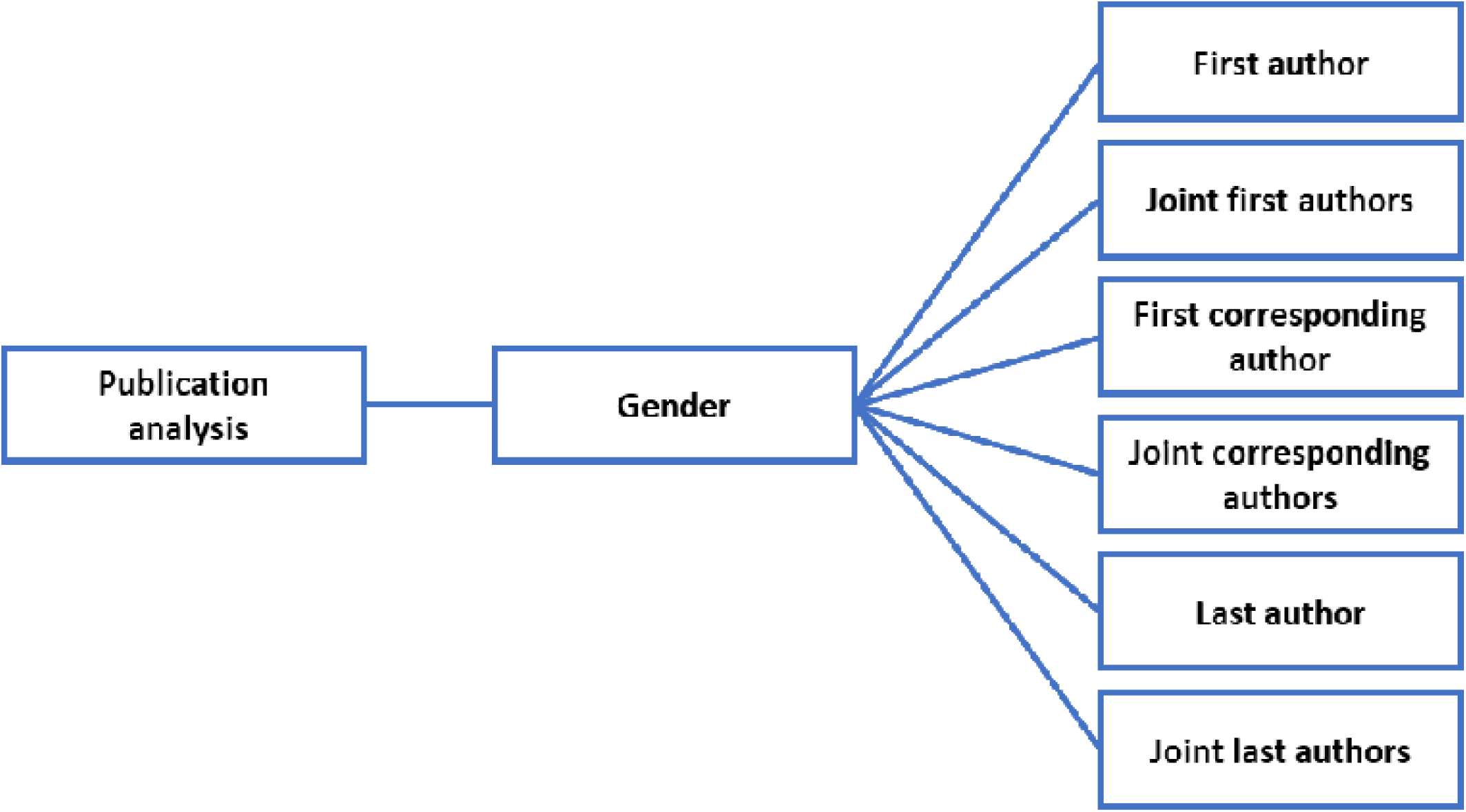
Publication analysis workflow.

First author was defined as the first-named author of the publication. Publications that consisted of single authors were categorised as first authors. We considered the first author to be the main intellectual contributor in the publication, in terms of study design, data collection and analysis, and manuscript writing. Joint first authors were defined as two or more authors who were named as equal contributors and mentioned as joint first authors of the publication. The first corresponding author was defined as the only author who was reported as a corresponding author in the publication and his/her contact details such as an institutional address and/or an email address were provided for correspondence in the publication. Joint corresponding authors were defined as two or more authors who were listed or marked as corresponding authors and their contact details were provided for correspondence in the publication. Last author was defined as the last-named author of a publication. The last author was considered to be a group leader or principal investigator who may have provided significant intellectual contribution or supervision of the research work as well as acquisition of research funding[17,35]. Joint last authors were defined as two or more authors who were named as equal contributors in the publication and their names were mentioned as joint last authors in the publication. A major confounding factor, for which we could not control, was the informal nature of the conventions for the sequence and role of authors[35]. Although conventions for scientific authorship are well established in biomedical sciences[36,37], they may vary between different research areas and even between different research groups within the same area.

### Determination of gender of authors

The gender of the authors was defined as a binary variable comprising either male or female categories, which were determined based on the first name of authors in all six categories of authorship included in the analysis. When the first names of authors were initialled in the publication or were difficult to associate with either male or female gender, further information was sought through searching their institutional webpages and online social network sites such as the LinkedIn and ResearchGate. We also used the Gender API (gender-api.com) when it was not possible to ascertain the gender of the authors by the above-mentioned sources. In addition, we contacted five authors directly via email to ascertain their gender. After completing data coding by two researchers (MJM and RD), to ensure the accuracy of data coding, 10% of the data were checked independently (CH). Consensus was achieved through discussion between the researchers on data fields that did not match the assigning of the gender of authors and types of authorship (Figure 1).

### Statistical analysis

Data were analysed using frequencies including counts and percentages. Chi-square tests were used for identifying statistically significant differences and associations between male and female authors in various categories of authorship. The level of significance was set at p < 0.05. Data were analysed using the IBM SPSS Statistics for Windows, Version 25.0 (Armonk, NY: IBM Corp.).

### Patient and public involvement statement

There was no patient or public involvement in the study design.

## RESULTS

### Type of research study

Table 1 presents an overview of the types of research studies by year. Clinical research studies (both trial and non-trial studies) comprised 39% (n=939), basic science research 27% (n=643) and a third of publications (34%, n=827) included other types of research, such as systematic reviews, reviews, research protocols, editorials, guidelines, opinions, comments, and meeting reports.

**Table 1.**
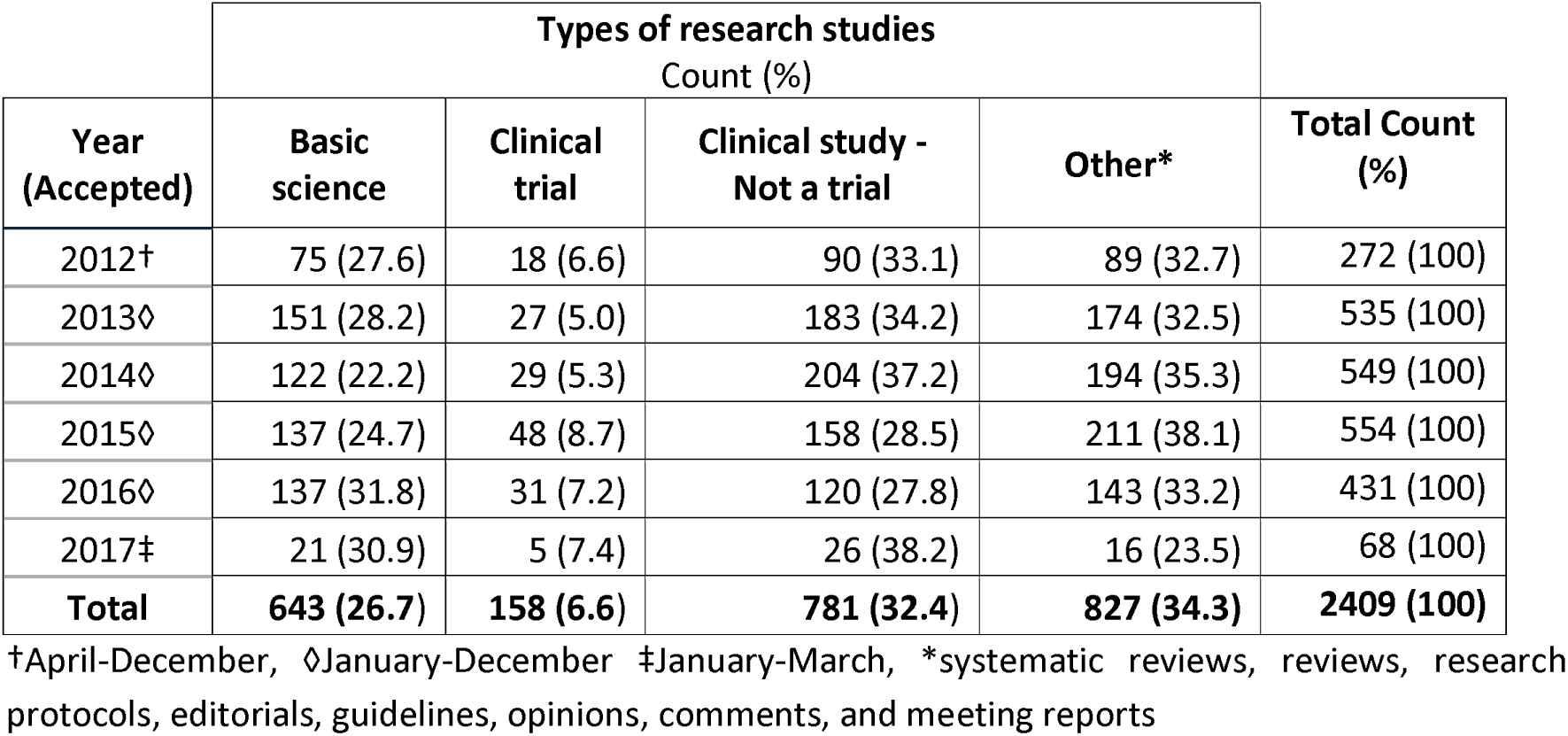
Number of publications by year of acceptance and types of research studies.

### Authorship type and Gender

Table 2 presents an overview of gender of authors by types of authorship. This highlights that male authors were statistically significant more likely to be first authors (59%, χ^2^ 972.938, P <0.001), first corresponding authors (66%, χ^2^ 242.970, P <0.001) and last authors (77%, χ^2^ 702.411, P <0.001)) (Table 2). Furthermore, analyses of joint authorship categories revealed that the proportion of ‘female only’ authors was statistically significantly lower than ‘male only’ authors in the joint corresponding authors (29%, χ^2^ 79.858, P<0.001) and joint last authors categories (10%, χ^2^ 56.550, P<0.001) (Table 2). However, in the joint first authors category, the proportion of ‘male and female’ as joint first authors (57%, χ^2^ 128.467, P <0.001) was statistically significantly higher than male only and female only first authors (Table 2).

**Table 2.** Authorship categories and gender of authors.

| Authorship type<br>Number of publications in the category | Gender of authors<br>Count (%) | | | Chi-Square Test<br>$\chi^2$ (p) |
| --- | --- | --- | --- | --- |
|  | Male only | Female only | Male and female |  |
| First author (n=2407) | 1413 (58.7) | 994 (41.3) | N/A * | 72.938 (<0.001) |
| First corresponding author (n=2371) | 1565 (66) | 806 (34) | N/A | 242.970 (<0.001) |
| Last author (n=2406) | 1853 (77) | 553 (23) | N/A | 702.411 (<0.001) |
| Joint first authors (n=458) | 127 (27.7) | 69 (15.1) | 262 (57.2) | 128.467 (<0.001) |
| Joint corresponding authors (n=169) | 107 (63.3) | 49 (29) | 13 (7.7) | 79.858 (<0.001) |
| Joint last authors (n=229) | 108 (47.2) | 23 (10) | 98 (42.8) | 56.550 (<0.001) |

\*N/A= not applicable.

### Gender of authors by type of research studies

Analysis of gender of authors by types of research studies (i.e. basic science, clinical trials, non-trial clinical studies and other research) showed that the proportions of male only authors were statistically significantly higher than the proportions of female only authors in three authorship categories: first authors (χ^2^ 8.606 (df 3), P = 0.035), first corresponding authors (χ^2^ 36.955 (df 9), P < 0.001) and last authors (χ^2^ 10.314 (df 3), P= 0.016). The analysis by type of research studies also revealed that there were no significant differences between the proportions of male only and female only authors in all three joint authorship categories: joint first authors (χ^2^ 5.549 (df 6), P = 0.476), joint corresponding authors (χ^2^ 9.021 (df 6), P = 0 .172) and joint last authors (χ^2^ 8.433 (df 6), P = 0 .208).

### Yearly trends in Authorship by gender

Figure 2 presents the yearly trends in scientific authorship by gender. In all authorship types and across all five years of publication, the proportion of male and female authors varied (Figure 2). The analysis showed women were significantly underrepresented across all years and authorship types. Interestingly, joint first authorship indicated a higher proportion of ‘male and female’ authors compared to ‘male only’ and ‘female only’ authors (Figure 2).

### Association between same gender across authorship categories

There was a statistically significant association between the same gender in first authorship and corresponding authorship categories (χ^2^ 775.425 (df 3), P < 0.001) and the first author and joint first authors (χ^2^ 138.849 (df 2), P < 0.001).

Furthermore, there were statistically significant associations between the same gender in the last author category with the same gender of first author(s) (χ ^2^ 33.742 (df 2), P < 0.001), corresponding author(s) (χ^2^ 540.774 (df 1), P < 0.001) and joint last authors (χ^2^ 91.291 (df 2), P < 0.001). However, there was no statistically significant association between the male and female last authors with the respective gender of joint first authors (χ^2^ 4.29 (df 2), P = 0.117).

## DISCUSSION

We retrospectively analysed the gender parity of authors in six categories of authorship of scientific publications that were published over a five-year period. Our analysis shows that the number of female authors were underrepresented across all six categories of authorship [10,38,39].

In the first author category the proportions of female authors and male authors were within the 40%-60% “gender balance zone”[6]. The greatest gender imbalance was observed in the last author category where ‘female only’ authors comprised only 23%. Nonetheless, this proportion is higher than other studies reporting similar analyses[11,16,24].

To the best of our knowledge, this study presents the first analysis of joint authorship in three categories. Secondly, it demonstrates underrepresentation in female only authors in six categories of scientific authorship[40]. Thirdly, the analysis highlights gender inequity with female underrepresentation in prestigious authorship positions compared to male in biomedical research. This is consistent with other fields including: epilepsy, lung cancer, dermatology, eating disorders and in medicine in general[17,19,41–43].

This study extends understanding of gender-based trends in scientific authorship (Figure2) by showing encouraging incremental changes in gender parity in authorship in a biomedical research setting. Previous research examined the gender gap in authorship within the medical literature reporting an upward trend for female first authors from 6% in 1970 to 29% in 2004 and female last authors from 4% in 1970 to 19% 2004. However, it was limited to US based institutions[12]. A similar UK based study covering the same period (i.e. 1970-2004) also showed upward trends for female first authors increasing from 11% in 1970 to 37% in 2004 and female last authors from 12% in 1970 to 17% in 2004[24]. In addition, a recent study by Filardo et al.[16] examined the prevalence of female first authorship of original research published in six high impact general medical journals between February 1994 and June 2014 revealed that the adjusted probability of an article having a female first authorship increased significantly from 27% in 1994 to 37% in 2014[16]. However, despite the proportion of female first authors varied greatly by journal, men were generally more likely to be first authors than women[16]. Compared to previous studies mentioned above, our study provides evidence of higher and increasing gender equity in the first authors, last authors and other four categories of scientific authorship in biomedical research (Table 2).

Our study identified a strong association between same gender and authorship types showing if the first author of a publication was male, it was highly likely that the first corresponding author of the same publication would also be male. Similarly, the likelihood of the first author being female was higher, if the first corresponding author was also female[44]. Likewise, there appeared to be a significant association of male and female last authors with the respective gender of first authors. Previous research has highlighted males and females were more likely to be first authors on papers if the last authors were of the same gender; however, these were not conducted in a translational research setting[23,45–47]. There was also a strong association of male and female last authors with the respective gender of corresponding authors[44].

However, due to the differences in gender equity between different research areas and medical specialties, where a centre-specific mix of research themes is likely to influence gender equity in scientific authorship, it is difficult to make direct comparisons across the literature.

Overall, our results build an important evidence base in biomedical research settings concerning gender parity and support the findings from previous studies where analysis of scientific authorship by gender has been used as an important marker of gender equity[12,24,48–50].

### Implications for policy and practice

While NIHR BRCs routinely collect bibliometric data on publications arising from the NIHR-funded research, and report to the NIHR (the funder), to the best of our knowledge, this data is not routinely analysed by gender. Our study supports the feasibility of using NIHR BRCs funded or supported research publications for analysing scientific authorship by gender. While retrospective analysis of the gender of authors in scientific publications is labour-intensive and has limitations, there is an opportunity to begin to track this prospectively. As more data becomes available, this would enable longitudinal analysis of scientific authorship by gender, which could be useful for tracking progress towards gender equity and related issues such as markers of achievement across all NIHR BRCs.

In addition, since the acceleration of women’s advancement and leadership in translational research is one of the stated objectives of the NIHR, investigating the extent of gender equity in scientific authorship may usefully inform strategies to accelerate women’s advancement and leadership in NIHR-funded research. Moreover, bibliometric analyses used by the NIHR to inform competition for NIHR funding may incorporate the gender dimension into the analysis, which could provide additional information on the competitiveness for NIHR funding[51,52].

## CONCLUSION

Our results show that while first authorship is within the 40%-60% gender balance zone, a greater gender disparity is prevalent in other types of scientific authorship in biomedical research. The proportion of female authors is significantly lower than the proportion of male authors in all six categories of authorship included in our analysis. This study also demonstrates the feasibility of analysing scientific authorship by gender, which could provide useful insight about gender equity in scientific publications, which may be a useful marker of achievement. Overall, our study demonstrates that it is feasible to analyse the available bibliometric data on publications arising from NIHR funding by gender and consider establishing processes for analysing gender equity in scientific authorship over time.

## Supporting information

Supplementary file 2 _Figure 2_Authorship trends by gender_400dpi

## Contributors

LDE conceived the study. RD and MJM coded the data. SGSS analysed the data. RD and SGSS drafted the manuscript. PVO contributed to the conception of the study and co-wrote parts of the manuscript. CRH and OC participated in data collection. VK, LRH, and AMB contributed to the conception of the study, facilitated access to the publications and coordinated the study. All authors read, contributed to and approved the final manuscript.

## Ethics

The University of Oxford Clinical Trials and Research Governance Team reviewed the study and deemed it exempt from full ethics review on the grounds that it falls outside of the Governance Arrangements for Research Ethics Committees (GAfREC), which stipulate which research studies are required to have ethics review. A wider programme of research on the activities of the NIHR Oxford Biomedical Research Centre from 2017 to 2022 received ethics clearance through the University of Oxford Central University Research Ethics Committee (R51801/ RE001), the Health Research Authority (IRAS ID 228049) and the Oxford University Hospitals NHS Foundation Trust Management (PID 12779).

## Acknowledgements

The authors wish to thank Professor Helen McShane for reviewing the draft manuscript and making comments and suggestions for improving it.

## Patient consent for publication

Not required.

## Data sharing statement

No additional data available.

## Additional files list

**Figure 1** Publication analysis workflow. The workflow shows the process of extracting data according to gender from six types of authorship.

**Figure 2** Yearly trends in scientific authorship by gender (male and female), April 2012 - March 2017. This plot represents the yearly variation of the proportion of male and female authors according to six types of authorship between the years of publication/acceptance from 2012 to 2017.

Supplementary File 1

