## Supplementary figures and images for "A retrospective analysis of gender parity in scientific authorship in a biomedical research centre"

### Supplementary file 2 _Figure 2_Authorship trends by gender_400dpi

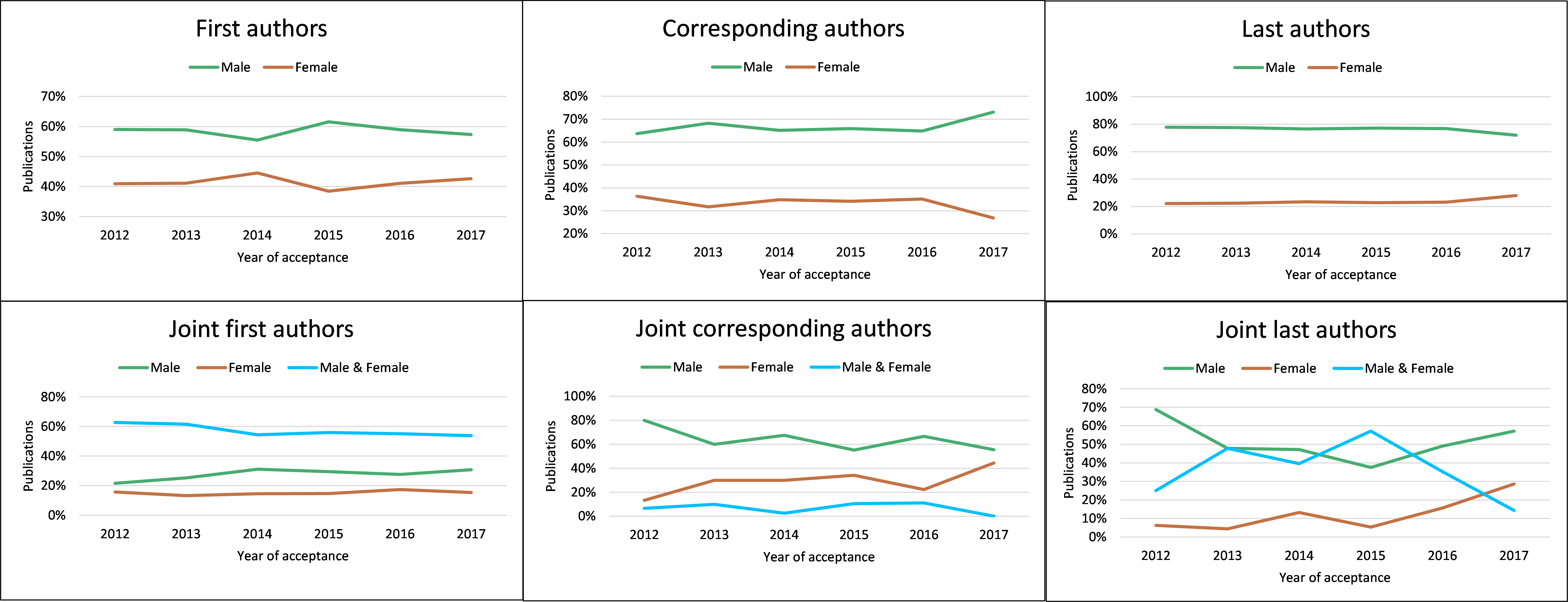
